# Integrative approaches for large-scale transcriptome-wide association studies

**DOI:** 10.1101/024083

**Authors:** Alexander Gusev, Arthur Ko, Huwenbo Shi, Gaurav Bhatia, Wonil Chung, Brenda W J H Penninx, Rick Jansen, Eco JC de Geus, Dorret I Boomsma, Fred A Wright, Patrick F Sullivan, Elina Nikkola, Marcus Alvarez, Mete Civelek, Aldonis J. Lusis, Terho Lehtimäki, Emma Raitoharju, Mika Kähönen, Ilkka Seppälä, Olli T. Raitakari, Johanna Kuusisto, Markku Laakso, Alkes L. Price, Päivi Pajukanta, Bogdan Pasaniuc

## Abstract

Many genetic variants influence complex traits by modulating gene expression, thus altering the abundance levels of one or multiple proteins. In this work we introduce a powerful strategy that integrates gene expression measurements with large-scale genome-wide association data to identify genes whose cis-regulated expression is associated to complex traits. We use a relatively small reference panel of individuals for which both genetic variation and gene expression have been measured to impute gene expression into large cohorts of individuals and identify expression-trait associations. We extend our methods to allow for indirect imputation of the expression-trait association from summary association statistics of large-scale GWAS^1-3^. We applied our approaches to expression data from blood and adipose tissue measured in ∼3,000 individuals overall. We then imputed gene expression into GWAS data from over 900,000 phenotype measurements^4-6^ to identify 69 novel genes significantly associated to obesity-related traits (BMI, lipids, and height). Many of the novel genes were associated with relevant phenotypes in the Hybrid Mouse Diversity Panel. Overall our results showcase the power of integrating genotype, gene expression and phenotype to gain insights into the genetic basis of complex traits.

## Introduction

Although a large proportion of variability in complex human traits is due to genetic variation, the mechanistic steps between genetic variation and trait are generally not understood^7^. Many genetic variants influence complex traits by modulating gene expression, thus altering the abundance levels of one or multiple proteins^8-12^. Such relationships between gene expression and trait could be investigated through association scans in individuals for which both measurements are available^8,13,14^. Unfortunately, studies that measure gene expression have been held back by specimen availability and cost, with the few published studies of gene expression and complex trait being orders of magnitude smaller than studies of trait alone. Consequently, many expression-trait associations cannot be detected, especially those with small effects. To mitigate the reduced power from small sample size, alternative approaches examined the overlap of genetic variants that impact gene expression (eQTLs) with trait-associated variants identified in large, independent genome-wide association studies (GWAS)^4,5,8,9,11-13,15^. However, this approach is likely to miss expression-trait associations of small effect.

We developed a new approach to identify genes whose expression is significantly associated to complex traits in individuals without directly measured expression levels (Methods). We leveraged a relatively small set of reference individuals for whom both gene expression and genetic variation (single nucleotide polymorphisms, SNPs) have been measured to impute the cis-genetic component of expression into a much larger set of phenotyped individuals from their SNP genotype data (Figure 1). We then correlated the imputed gene expression to the trait to perform a transcriptome-wide association study (TWAS) and identify significant expression-trait associations (Methods). A critical limitation is that large-scale GWAS data are typically only publicly available at the level of summary association statistics (e.g. individual SNP effect sizes)^1-3^. We extended our approach to impute expression-trait association statistics at the summary statistic level (Methods). This allowed us to increase the effective sample size for expression-trait association testing to hundreds of thousands of individuals. By focusing only on the genetic component of expression, we avoid instances of expression-trait associations that are not a consequence of genetic variation but are driven by variation in trait (Figure 2). Our approach can be conceptualized as a test for significant cis-genetic correlation between expression and trait (Methods).

**Figure 1:**
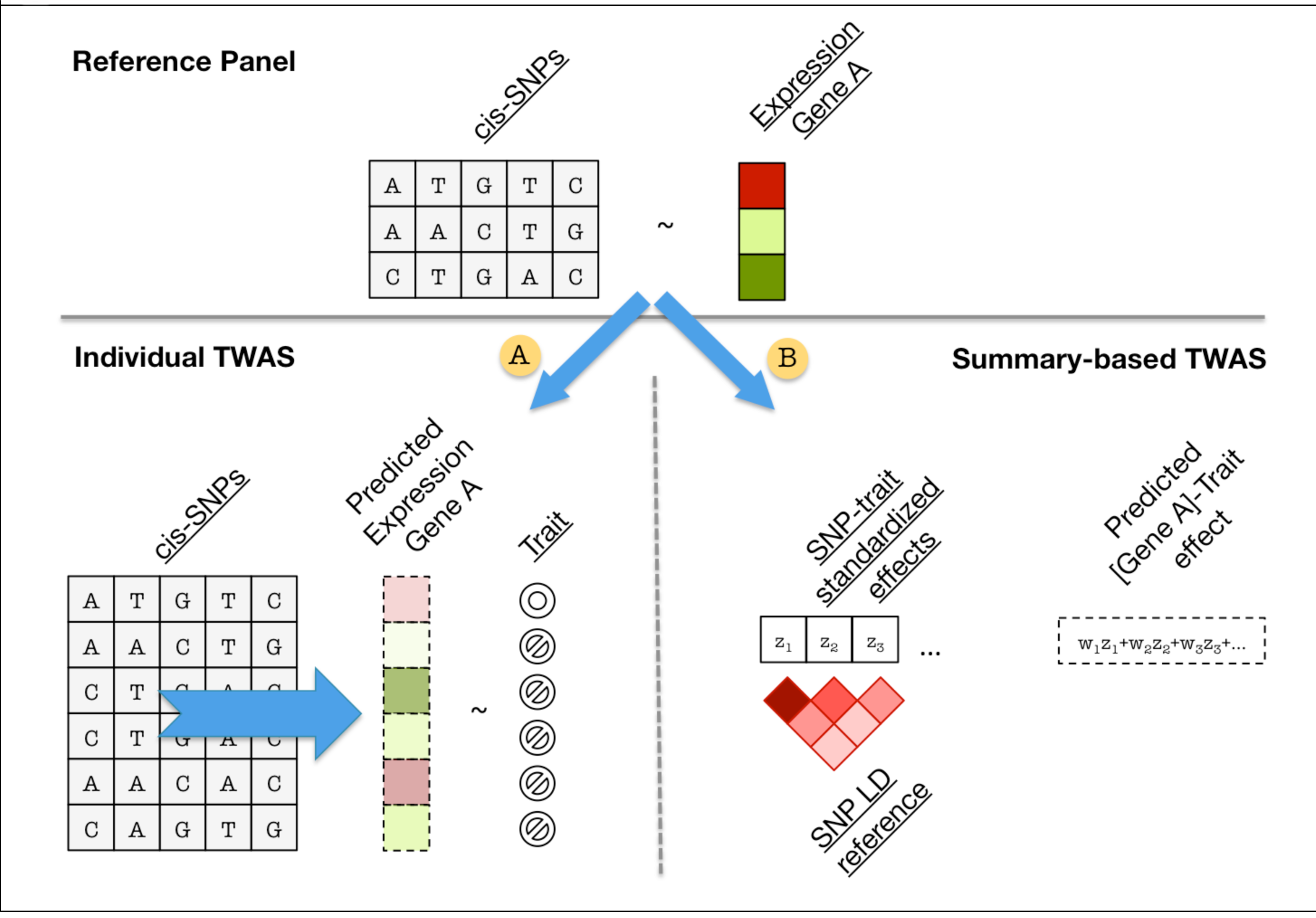
**Overview of methods.** Cartoon representation of TWAS approach. In the reference panel (top) estimate gene expression effect-sizes: directly (i.e. eQTL); modeling LD (BLUP); or modeling LD and effect-sizes (BSLMM). A: Predict expression directly into genotyped samples using effect-sizes from the reference panel and measure association between predicted expression and trait. B: Indirectly estimate association between predicted expression and trait as weighted linear combination of SNP-trait standardized effect sizes while accounting for LD among SNPs.

**Figure 2:**
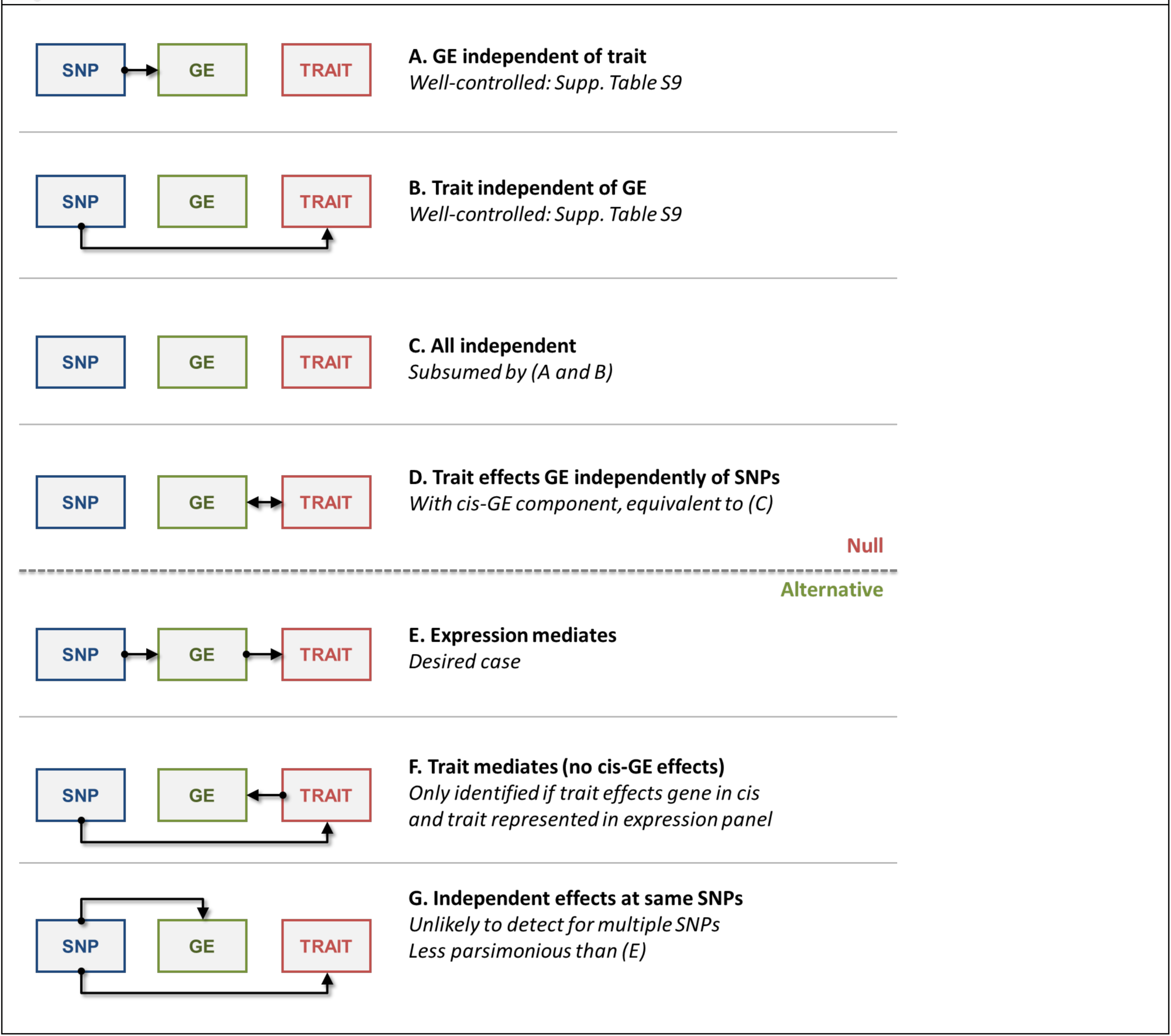
**Modes of expression causality.** Diagrams are shown for the possible modes of causality for the relationship between genetic markers (SNP, blue), gene expression (GE, green), and trait (red). A-D describes scenarios that would be considered null by the TWAS model; E-G describes scenarios that could be identified as significant.

We applied our approaches to expression data from blood and adipose tissue measured in ∼3,000 individuals overall. Through extensive simulations and real data analyses we show that our proposed approach increases performance over standard GWAS or eQTL-guided GWAS. Furthermore, we reanalyzed a 2010 lipid GWAS^16^ to find 25 new expression-trait associations in that data. 19 out of 25 contained genome-wide significant SNPs in the more recent and expanded lipids study^4^ thus showcasing the power of our approach to find robust associations. We imputed gene expression into GWAS data from over 900,000 phenotype measurements^4-6^ to identify 69 novel genes significantly associated to obesity-related traits (BMI, lipids, and height). Many of the novel genes were associated with relevant phenotypes in the Hybrid Mouse Diversity Panel. Overall our results showcase the power of integrating genotype, gene expression and phenotype to gain insights into the genetic basis of complex traits.

## Results

### Overview of the methods

We integrate gene expression and GWAS data using imputed gene expression from SNP data. Briefly, we use a reference panel of gene expression and SNP data to build a statistical model for predicting the cis-genetic component of expression. The imputed expression can be viewed as a linear model of SNP data with weights based on the correlation between SNPs and gene expression in the training data while accounting for linkage disequilibrium (LD) among SNPs. To capitalize on the largest GWAS to date (typically available only at the summary level), we extended our approach to impute the expression-trait association statistics directly from GWAS summary statistics (Methods). In contrast to expression imputation from individual-level data, imputation of expression-trait association from GWAS summary statistics can exploit publically available data from hundreds of thousands of samples. Linear predictors naturally extend to indirect imputation of the standardized effect of the cis-genetic component on the trait starting from only the GWAS association statistics^1-3^. Similarly to individual level imputation, the predicted effect of the expression on the trait can be viewed as a linear combination of the effects of each SNP on the trait with weights estimated from reference panels that contain both SNP and gene expression measurements (see Methods).

### SNP-heritability of gene expression

To investigate the potential utility of a transcriptome-wide association (TWAS) based on imputed gene expression we first estimated the cis- (1Mb window around the gene) and trans-(rest of the genome) SNP-heritability (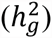) for each gene in our data^17,18^. These metrics quantify the maximum possible accuracy (in terms of R^2^) of a linear predictor from the corresponding set of SNPs^19,20^ (Methods). We used 3,234 individuals for whom genome-wide SNP data and expression measurements were available: 573 individuals with RNA-seq measured in adipose tissue from the METSIM (METabolic Syndrome In Men) study^41,42^; 1,414 individuals with array-based expression measured in whole blood from the Young Finns Study (YFS)^21,22^; and 1,247 unrelated individuals with array-based expression measured in peripheral blood (NTR, Netherlands Twin Registry)^23^ (Methods, Table S1). All expression measurements were adjusted for batch confounders, and array probes were merged into a single expression value for each gene where possible (Methods). Consistent with previous work^23,24^, we observed significantly non-zero estimates of heritability across all three studies, with mean 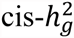 ranging from 0.01-0.07 and mean 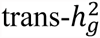 ranging from 0.04-0.06 in genes where estimates converged (Figure S3, Table S1). Although we observed large differences in the average 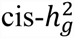 estimates between the two blood cohorts, the estimates were strongly correlated across genes (Pearson *ρ*=0.47 for YFS-NTR, as compared to *ρ*=0.15 and *ρ*=0.26 for METSIM-NTR and METSIM-YFS respectively). This is consistent with a common but not identical genetic architecture. The 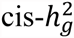 was significantly non-zero for 6,924 genes after accounting for multiple hypotheses (1,985 for METSIM, 3,836 for YFS, and 1,103 for NTR) (Figure S3) whereas current sample sizes where too small to detect individually significant trans heritable genes. As expected, we also observed a high overlap of genes with significant 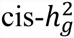 across cohorts (Table S15). We focused subsequent analyses on the 6,924 cis-heritable genes as such genes are typically enriched for trait associations^6,9,13,23-28^.

### TWAS performance in simulations

We evaluated whether the expressions of the 6,924 highly heritable genes could be accurately imputed from cis-SNP genotype data alone in these three cohorts. In each tissue, we used cross-validation to compare predictions from the best cis-eQTL to those from all SNPs at the locus either in a best linear unbiased predictor (BLUP) or in a Bayesian model^29,30^ (Methods). On average, the Bayesian linear mixed model (BSLMM)^30^, which uses all cis-SNPs and estimates the underlying effect-size distribution, attained the best performance with a 32% gain in prediction R^2^ over a prediction computed using only the top cis-eQTL (Figure 4, Figure S1). The BSLMM exhibited a long tail of increased accuracy, more than doubling the prediction R^2^ for 25% of genes (Figure S2). In contrast to complex traits where hundreds of thousands of training samples are required for accurate prediction^31,32^, a substantial portion of variance in expression can be predicted at current sample sizes due to the much smaller number of independent SNPs in the cis region^20^. Furthermore, larger training sizes will continue to increase the total number of genes that can be accurately predicted (Figure 3).

**Figure 3:**
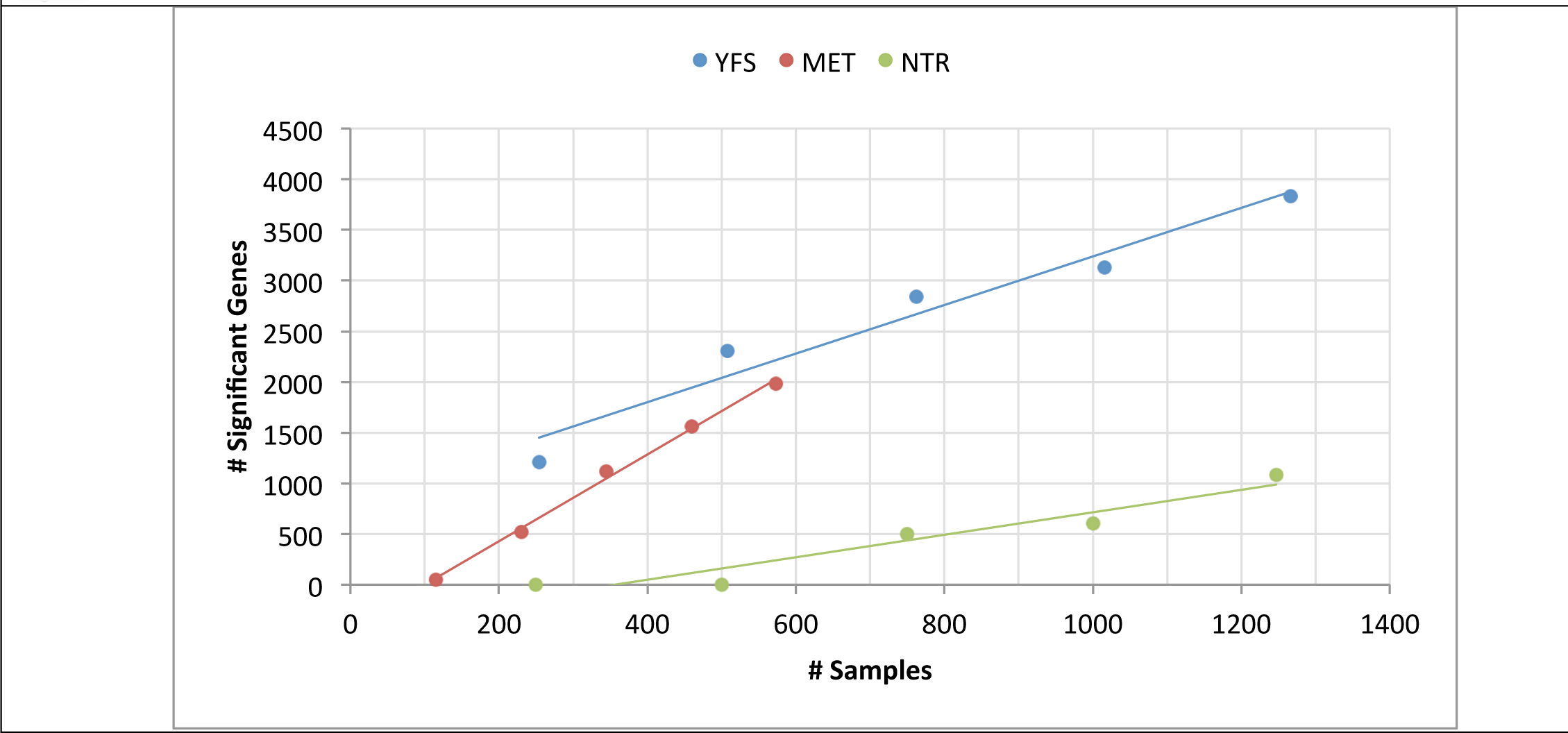
**Number of genes with significant cis-heritability observed at varying sample sizes.** The number of genes with significant cis-heritability was estimated by down-sampling each cohort (YFS, METSIM, and NTR/Wright et al.) into quintiles.

**Figure 4:**
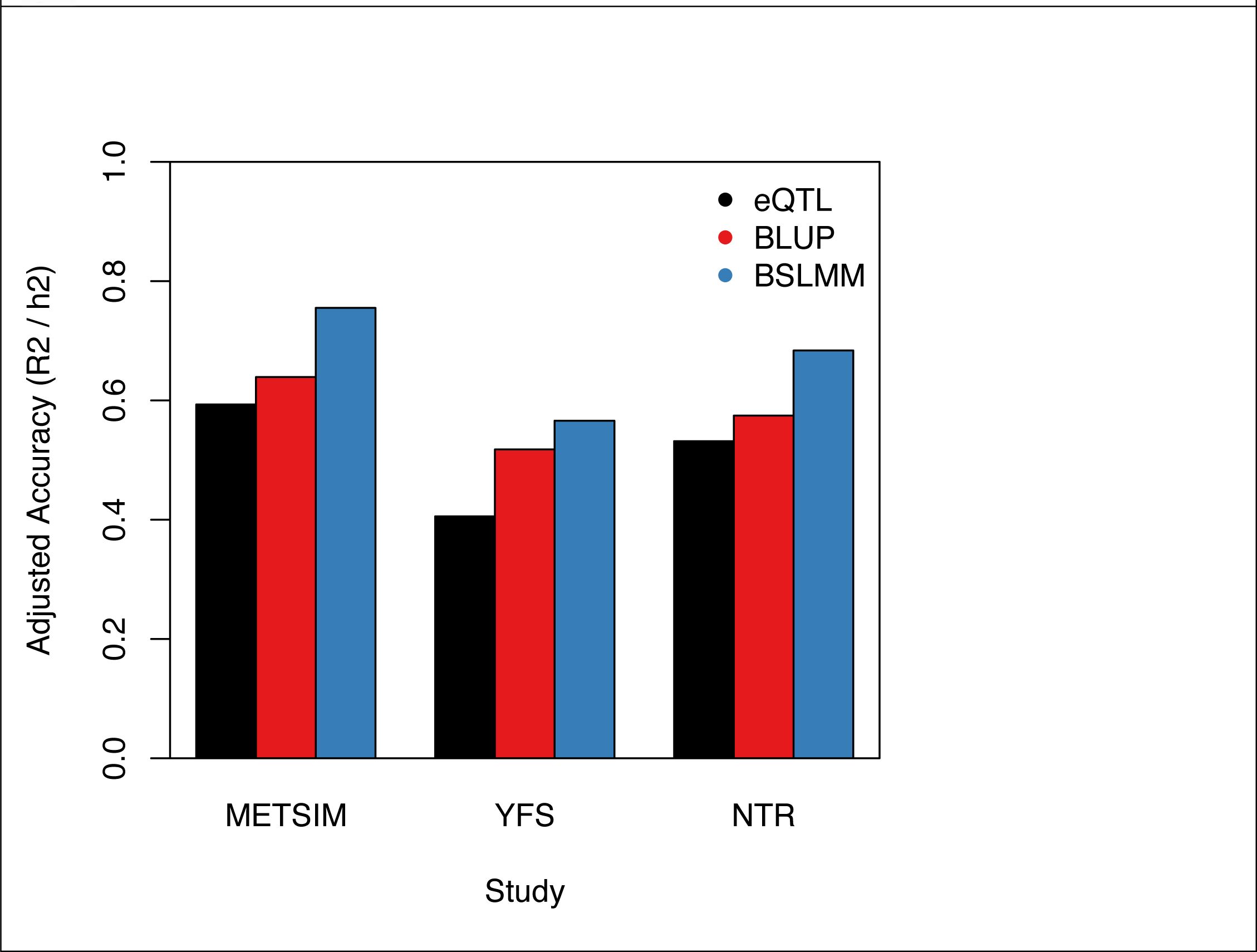
**Accuracy of direct expression imputation algorithms.** Adjusted accuracy was estimated using cross-validation R^2 between prediction and true expression, and normalized by corresponding cis-h2g. Bars show mean estimate across three cohorts and three methods: eQTL – single best cis-eQTL in the locus; BLUP using all SNPs in the locus; BSLMM using all SNPs in the locus and non-infinitesimal priors.

Next, we focused on evaluating the power of the TWAS approach to detect significant expression-trait associations using GWAS summary data from complex traits (equivalent to TWAS from individual level data; see Methods, Supplementary Figure S13). For comparison, we also measured power to detect significant SNP-trait associations through standard GWAS (testing each SNP individually) and eQTL-based GWAS (eGWAS, where the best eQTL in each gene is the only variant tested for association to trait), with all three tests corrected for their genome-wide test burdens. Using real genotype data, we simulated a causal SNP-expression-trait model with realistic effect-sizes and measured the power of each strategy to identify genome-wide significant variants (accounting for 1 million SNPs for GWAS and 15,000 expressed genes using family-wise error rate control). Over many diverse disease architectures (Methods) TWAS substantially increased power when the expression-causing variants were un-typed or poorly tagged by an individual SNP (Figures 5, S8, S9, S10, S11, S12, S14). The greatest power gains were observed in the case of multiple causal variants: 92% power for TWAS compared to 18% and 25% for GWAS and eGWAS. This scenario would correspond to expression caused by allelic heterogeneity^9,33,34^, or “apparent” heterogeneity at common variants (due to tagging of unobserved causal variant)^35^. TWAS was comparable to other approaches when a single causal variant was directly typed, in which case combining the effects of neighboring SNPs does not add signal. Under the null where expression was completely independent of phenotype (with either being heritable, Figure 2A-D), the TWAS false positive rate was well controlled (Table S9). As expected, all methods were confounded in the case where the same causal variants had independent effects on trait and expression (Figure 2F-G; Supplementary Figure S9, S14).

We also compared TWAS to a recently proposed method, COLOC^36^, for evaluating co-localization of gene expression at known GWAS risk loci. After matching the false-discovery rate of the two methods in simulations (Methods), we observed that TWAS and COLOC had similar power under the single typed causal variant scenario (with slightly lower COLOC power at small GWAS sizes), but that TWAS has superior performance when the causal variant was un-typed or in the presence of allelic heterogeneity (Supplementary Figure S11). This is likely due the fact that TWAS explicitly models LD to better capture the un-typed variants. Though COLOC has lower power to detect new loci, it offers the advantage of testing specific biological hypotheses and may be useful in evaluating known associations.

Finally, we investigated the effect of the expression reference panel size on performance of TWAS (see Supplementary Figure S10). In general, we observe that the TWAS approach always outperforms eGWAS when multiple variants are causal. Interestingly, power for either approach does not increase substantially beyond 1,000 expression samples, suggesting that the expression panels analyzed in this manuscript nearly saturate the available power. Although these simulation results come with caveats (e.g. standard assumptions of additive effects and normal residuals), they suggest that the main benefit of larger reference panels of expression data is in increasing the total number of significantly cis-heritable genes available for imputation (Figure 3).

### TWAS performance in real phenotype data

We assessed the performance of TWAS in identifying trait-effecting genes at the 697 known GWAS risk loci for height^6^ using the YFS expression samples for which height was also measured. At each locus, we considered three strategies for selecting a single causal gene: 1) the gene nearest to the most significantly associated SNP; 2) the gene for which the index SNP is the strongest eQTL in the training data; 3) the most significant TWAS gene. For each strategy, we then constructed a risk-score using the genetic value of expression for the selected genes and correlated the risk score with height measurements in the YFS individuals (an independent sample from the original height GWAS, see Methods). The R^2^ between the risk score and height was 0.038 (nearest); 0.031 (eQTL); and 0.054 (TWAS); with TWAS significantly higher than the others in a joint model (Table S3). Although functional validation is required, these results suggest that TWAS can be used to prioritize putative risk genes at known GWAS loci. Across all known loci from four GWAS, 61% of genome-wide significant loci (defined as lead SNP +/- 500kb) overlapped at least one significant cis-h2g gene, and 17% contained at least one significant TWAS association (Table S16). This suggests a substantial correlation of cis-expression and trait-effecting SNPs even at current power. Out of the 372(33) genes significant for height (BMI) in our analysis, 64 (1) are also reported in the original studies. We note that sensitivity may be low either due to a causal mechanism that does not involve cis-expression of the tested genes, or low power to identify and detect all cis-heritable genes at the locus.

Next, we employed TWAS to identify novel expression-trait associations using summary association statistics from a 2010 lipid GWAS^16^ (∼100,000 samples), i.e. associations that did not overlap genome-wide significant SNPs in that study. We used all three studies (METSIM, YFS, and NTR; Table S1) as separate SNP-expression training panels. We then looked for genome-wide significant SNPs at these loci in the larger 2013 lipid GWAS^4^ (expanded to ∼189,000 samples). We identified 25 such expression-trait associations in the 2010 study (Table S4), of which 19/25 contained genome-wide significant SNPs in the 2013 study (P=1×10^−24^ by hypergeometric test, Methods) and 24/25 contained a more significant SNP (P=1×10^−04^), a highly significant validation of the predicted loci. The validation remained significant after conservatively accounting for sample overlap across the studies (binomial P=3×10^−16^; Methods, Table S4). As a sanity check, we compared direct and summary-level TWAS in the METSIM cohort, and found the two sets of imputed expression-trait Z-scores to be nearly identical, with summary-level TWAS slightly under-estimating the effect (Pearson *ρ*=0.96, Figure S5). Overall, we find the TWAS approach to be highly predictive of robust phenotypic associations.

### TWAS identifies novel expression-trait associations for obesity related traits

Having established the utility of TWAS, we applied the approach to identify novel expression-trait associations using summary data from three recent GWAS over more than 900,000 phenotype measurements: lipid measures (high-density lipoproteins [HDL] cholesterol, low-density lipoprotein [LDL] cholesterol, total cholesterol [TC], and triglycerides [TG])^4^; height^6^; and BMI^5^. Significantly cis-heritable genes across the three expression data sets were tested individually (6,924 tests) and together in an omnibus test that accounts for predictor correlation (1,075 tests; Methods), and we conservatively corrected for the 8,000 total tests performed for each trait. Overall, we identified 665 significant gene-trait associations (Table S5). Of these, 69 gene-trait associations did not overlap a genome-wide significant SNP in the corresponding GWAS, residing in 60 physically non-overlapping cis-loci (Table 1, Table S6). Averaging over the novel genes, the TWAS explained 1.5x more phenotypic variance than the strongest eQTL SNP for the same gene, and was more significant for 88% of the tests (though this may include some winner’s curse). Our previous simulations suggest that the substantial gain over testing the cis-eQTL is an indication of pervasive allelic heterogeneity^37^ at these loci. Our results are consistent with a model of causality where these genes harbor inherited causal variants that modulate expression, which in turn has a complex effect on the cell and downstream impact on complex traits^5^.

We further sought to quantify the significance of the expression-trait associations conditional on the SNP-trait effects at the locus with a permutation test (Methods). Comparing to this null assesses how much signal is added by the true expression given the specific architecture of the locus. Of the 69 genes, this permutation test was significant for 54 (after accounting for 69 tests). After excluding these individually significant genes, the P-values were still substantially elevated with λ_GC_ of 19 (ratio of median *χ*^2^ to the expected null). For these 54 genes, we can confidently conclude that integration of expression data significantly refined the association to trait.

For all expression-trait associations, including those that were not genome-wide significant, we estimated the aggregate contribution to phenotype using two heritability models. As has been shown previously, there is a relationship between the mean χ^2^ statistic and total variance in trait explained by the associated markers under a polygenic architecture (Methods,^29,30^). Leveraging this relationship, we estimated the variance in trait explained by all METSIM+YFS imputed genes 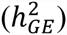 to be 3.4% averaged over six traits (Table S7). We assumed independence between the two cohorts, and did not include the NTR genes because of its strong correlation with YFS. Height had the most variance attributable to the heritable genes at 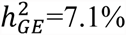. These combined estimates were consistently higher than a corresponding analysis using predictions from permuted expression (Table S7). For the four traits with individual-level genotype and phenotype data in the METSIM (BMI, TG, WHR, INS), we estimated 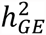 directly using variance-components over the imputed expression values (Methods). On average, all significantly heritable genes in adipose + blood explained 4-6% of the trait variance (16-19% of the total trait 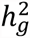), and were largely orthogonal between the two predictions (Table S8). The imputed expression consistently explained more trait variance than the best cis-eQTL in each gene and did not strongly depend on the cis-window size (Table S9).

### Re-evaluation using other expression cohorts

To replicate the 69 novel expression-trait associations, we re-evaluated the GWAS summary statistics with expression data from two external studies: eQTLs from ∼900 samples in the MuTHER study^24^ of fat, LCL, and skin cells; and separate eQTL data from a meta-analysis of 8,086 samples across multiple tissues from ref.^11^ (Methods). We note that these expression studies only consist of summary-level associations, and are expected to be much noisier as reference. In the relatively smaller MuTHER sample, 20 out of 55 available genes replicated significantly in at least one tissue (after accounting for 55 tests, Table S6). This is substantial given the apparent heterogeneity between cohorts we previously observed (Methods). Importantly, the correlations between discovery and replication Z-scores were strongest for associations found in the corresponding tissue (*ρ*=0.60, P=1.5×10^−05^ for blood/LCL; *ρ*=0.66, P=0.05 for adipose, Table S10); a significant aggregate replication and further evidence for the tissue-specific nature of our findings. Using the larger, but heterogeneous, training sample from ref.^11^, 24 out of 37 available genes replicated significantly (Table S6). Although these replications are not strictly independent (they use the same GWAS data), they demonstrate that many of the novel loci are consistently significant across diverse expression cohorts.

### Functional analysis of the novel associations

To better understand their functional consequences, we evaluated the 69 novel genes in the Hybrid Mouse Diversity Panel (HMDP) for correlation with multiple obesity-related traits. This panel includes 100 inbred mice strains with extensive collection of obesity-related phenotypes from ∼12,000 genes. Of the 69 novel TWAS genes previously identified, 40 were present in the panel and could be evaluated for effect on phenotype. Of these, 26 were significantly associated with at least one obesity-related trait (after accounting for genes tested) and 14 remained significant after accounting for 36 phenotypes tested (very conservatively assuming the phenotypes were independent) (Table S11). 77% of the genes with an association were associated with multiple phenotypes. For example, expression of *Ftsj3* was significantly correlated with fat mass, glucose-to-insulin ratio, and body weight in both liver and adipose tissue, with R^2^ ranging from 0.20-0.28. Another candidate, *Iih4*, was significantly correlated with LDL and TC levels in liver. In humans, this gene is also linked to hypercholesterolemia in OMIM and was previously associated with BMI in East Asians^38^. Due to complex correlation of phenotypes, it is difficult to assess whether this gene set is significant in aggregate and genes in the HMDP are typically expected to have strong effects. We could not perform enough random selections of genes to establish significance for this set. However, we consider the 26 individually significant genes to be fruitful targets for follow-up studies.

The BMI and height GWAS evaluated functional enrichment at identified loci, and we performed similar analyses for the novel genes that we identified. We tested the 10 novel BMI genes and 33 novel height genes for tissue-specific enrichment using DEPICT^39^, a method based on large-scale gene co-expression analyses, following the protocol of the original GWAS studies^5,6^. Analysis of BMI identified significant enrichment for hypothalamus and neurosecretory systems (P=2.6×10^−4^, significant at FDR<5%). This enrichment is consistent with the landmark finding in the original study^5^ showing enrichment in these and other central nervous system tissues. Notably, we recapitulated this result using only novel loci that did not overlap any genome-wide significant SNPs. In analysis of height, DEPICT did not identify any tissue-specific enrichment.

### Discussion

In this work we proposed methods that integrate genetic and transcriptional variation to identify genes with expression associated to complex traits. Using imputed gene expression to guide GWAS has three potential advantages. First, the gene is a more interpretable biological unit than an associated locus, which often contains multiple significant SNPs in LD that may not lie in genes and/or tag variants in multiple genes. Second, the lower total number of genes (or cis-heritable genes) means the multiple-testing burden is substantially reduced relative to all SNPs. Lastly, combining cis-SNPs into a single predictor may capture heterogeneous signal better than individual SNPs or cis-eQTLs. Focusing the prediction on the genetic component of expression also avoids confounding from environmental differences caused by the trait that may impact expression. Our approach builds upon the wealth of GWAS data in massive cohorts to directly implicate the gene-based mechanisms underlying complex traits. Our proposed method shares conceptual similarities with 2-sample Mendelian randomization approaches that aim to identify causal relations between traits using genetic variation predictions as a randomizer^40-42^. However, while Mendelian randomization is intended to quantify the total causal effect (in this case of expression on trait), our method has the less strict goal of identifying a significant genetic correlation (i.e. associations) and can operate on summary GWAS data. Importantly, our approach maintains the attractive feature of not being confounded by effects on expression and trait that are independent of the SNPs.

Unlike current methods, which focus on individually significant eQTL and SNP associations^4,5,8,9,11,13,25,28^, our approach captures the full cis-SNP signal and does not require any individual marker to be significant. This is underscored by the fact that the TWAS estimate substantially outperformed its cis-eQTL analog both in predicting expression and in explaining trait. Our simulations show that the imputation approach is especially effective when multiple variants influence expression (which in turn influences trait). The large number of novel associations identified in real data supports this phenomenon and suggests that it may be a strong contributor to common phenotypes^43^. In these cases, our approach can be seen as complementary to GWAS by identifying expression-trait associations that are not well explained by individual tagging SNPs. Future work could leverage the difference in performances of TWAS and GWAS to detect loci with allelic heterogeneity. We note that it is still possible for some loci to have an independent SNP-phenotype and SNP-expression association driven by the same underlying variant though we consider this to be an infrequent biological model.

We conclude with several limitations of our proposed approach. First, disease-impacting variants that are independent of cis-expression – in general or in the training cohort – will not be identified. Second, as with any prediction, the number of genes that can be accurately imputed is still limited by the training cohort size and the quality of the training data. In particular, we found that prediction accuracy did not correspond to theoretical expectations and is likely driven by data quality. The impact of these weaknesses could be better quantified as expression from larger sample sizes and a more diverse set of tissues becomes available. Although in this work we utilized both microarray and RNA-seq as measure of gene expression thus showcasing the applicability of our approach to diverse data sets, the accuracy of our method intrinsically depends on the quality of the expression measurements. For the associated genes, it remains possible that the effect is actually mediated by phenotype (i.e. SNP – phenotype – cis-expression, Figure 2F). We attempted to quantify this in the YFS data by conditioning the heritability analyses on all the evaluated phenotypes (height, BMI, and lipids) but observed no significant change at individual genes or in the mean cis-h2g. These results suggest that confounding from phenotype does not substantially affect the tested cis expression, though at the current sample size we cannot completely rule out such confounders for individual genes. An alternative confounder arises from independent effects on phenotype and expression at the same SNP/tag (Figure 2G, Methods). Such instances could be indistinguishable from the desired causal model (Methods) without analyzing individual-level data, though we believe they are still biologically interesting cases of co-localization. Both types of confounding could potentially be quantified by training the SNP-expression relationships in control individuals where phenotype is fixed, or by interrogating the gene experimentally. Lastly, the summary-based TWAS cannot account for rare variants that are poorly captured by the LD reference panel, or optimally capture non-linear relationships between SNPs and expression. Additional sources of information could potentially be incorporated to improve the prediction, including significant trans-associations^11,27^; allele-specific expression^44,45^; splice-QTLs effecting individual exons^10^; haplotype effects; and SNP-specific functional priors^19,46,47^.

**Figure 5:**
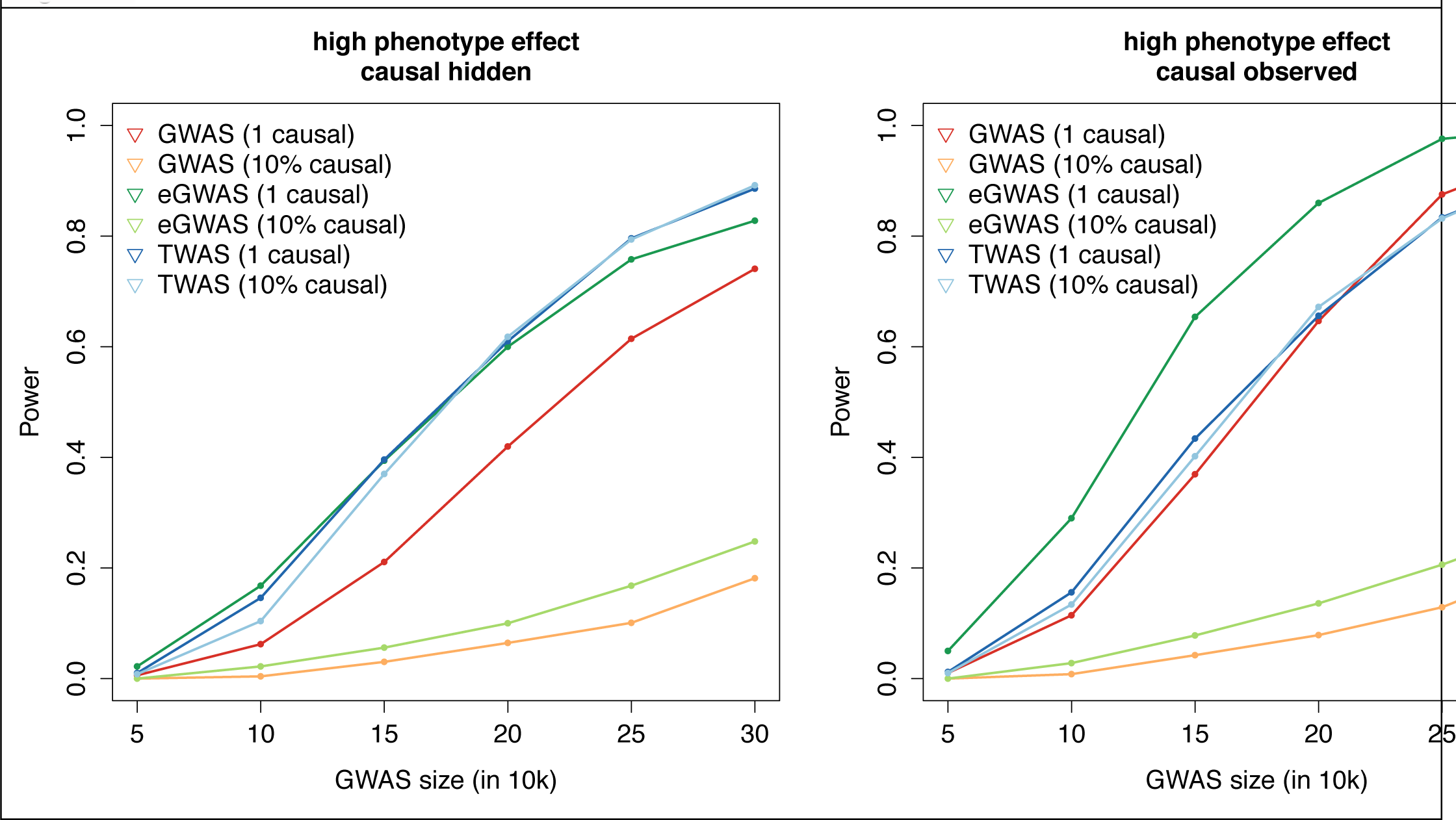
**Power of summary-based expression imputation algorithms.** Realistic disease architectures were simulated and power to detect a genome-wide significant association evaluated across three methods (accounting for 15,000 eGWAS/TWAS tests, and 1,000,000 GWAS tests). Colors correspond number of causal variants simulated and methods used: GWAS where every SNP in the locus is tested; eGWAS where only the best cis-eQTL is tested; and TWAS computed using summary-statistics. Expression reference panel was fixed at 1,000 out-of-sample individuals and simulated GWAS sample size designated by x-axis. Power was computed as the fraction of 500 simulations where significant association was identified.

## Methods

### METSIM and YFS cohorts

In this study, we included 11,484 participants from two Finnish population cohorts, the METabolic Syndrome in Men (METSIM, n=10,197)^48,49^ and the Young Finns Study (YFS, n=1,414)^21,22^. Both studies were approved by the local ethics committees and all participants gave an informed consent. The METSIM participants were all male with a median age of 57 years (range: 45-74 years) recruited at the University of Eastern Finland and Kuopio University Hospital, Kuopio, Finland. Whole blood was collected from all individuals for genotyping and biochemical measurements. Additionally, 1,400 randomly selected individuals from the 10,197 METSIM participants underwent a subcutaneous abdominal adipose biopsy of which 600 RNA samples were analyzed using RNA-seq. Traits BMI, TG, WHR, and INS were inverse rank transformed and adjusted for age and age-square. INS was additionally adjusted for T1D and T2D. There was little correlation between genome-wide SNP principal components and phenotype. A strictly unrelated subset of 5,501 individuals was computed by removing one of any individual with off-diagonal GRM entries >0.05 (with priority given to individuals that had expression measured). This procedure guaranteed no relatedness between the training set and the samples without expression.

YFS participants were originally recruited from five regions in Finland: Helsinki, Kuopio, Oulu, Tampere, and Turku. We collected whole blood into PAXgene tubes from all individuals for genotyping, RNA microarray assay, and biochemical measurements. Samples from 1,414 individuals (638 men with a median age of 43 years and 776 women with a median age of 43) with gene expression, phenotype, and genotype data available were included in the blood expression analysis. Traits height, BMI, TG, TC, HDL, and LDL were inverse rank transformed and adjusted for age, age-square, and sex. TC was also adjusted for Statin intake.

The biochemical lipid, glucose, and other clinical and metabolic measurements of METSIM and YFS were performed as described previously^21,48,50^.

### METSIM RNA-Seq data

We prepared and sequenced mRNA samples isolated from subcutaneous adipose tissue using Illumina TrueSeq RNA Prep Kit and the Illumina Hiseq 2000 platform to generate 50bp long paired-end reads. Reads were aligned to the Human reference genome, HG19, using the aligner STAR^51^, allowing up to 4 mismatches for each read-pair. Transcript quantification was calculated as reads per kilobase per million (RPKM) using Flux Capacitor^45^ based on transcript and gene definitions from the Gencode ver.18 annotation. The gene quantification is the sum of all transcripts of a gene. We applied Anscombe transformation to the RPKM values for variance stabilization followed by PEER^52^ correction to remove technical biases. The PEER-corrected gene quantification was then inverse rank transformed to a normal distribution to eliminate effect from outliers.

### YFS microarray data analysis

We measured mRNA expression in whole blood of the YFS cohort using the Illumina HumanHT-12 version 4 Expression BeadChip. Probe density data were exported from GenomeStudio and analyzed in R using Bioconductor packages. We normalized probe density data using control probes with the neqc function from the limma package implemented in R^53^. To account for technical artifacts, we first log2 transformed the normalized density and adjusted for 20 potential confounding factors using PEER^52^. The final adjusted probe density was inverse rank transformed to approximate normality in order to minimize the effect of outliers. Probes which contained a SNP in the 1000 Genomes were removed.

### NTR expression array data

Data from the NTR was processed as described in the original paper^23^, followed by removal of any individuals with GRM values > 0.05. For genes with multiple probes, total gene expression was measured as the sum across all probes followed by standardization to unit variance (rank normalization had no substantial effect). Probes which contained a SNP in the 1000 Genomes were removed. Principal components and batch were included as covariates in all analyses.

### Heritability estimation with individual data

Cis and trans variance components were estimated using the REML algorithm implemented in GCTA^18^. As in previous studies, estimates were allowed to converge outside the expected 0-1 bound on variance to achieve unbiased mean estimates across all genes^23^. Standard error across gene sets was estimated by dividing the observed standard deviation by the square root of the number of genes that converged (this will lead to underestimation due to correlated genes, but is presented for completeness). Genome-wide 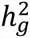 for the four traits in the GWAS cohort was estimated with GCTA from a single relatedness matrix constructed over all post-QC SNPs in the strictly unrelated individuals. For estimating expression-wide 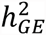, each predicted expression value was standardized to mean=0 and variance=1, and sample covariance across these values used to define the relatedness matrix. The 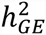 was then estimated from this component with GCTA, with P-values for difference from zero computed using a likelihood ratio test. 20 principal components (PCs) were always included as fixed-effects to account for ancestry. Genetic correlation between traits in the GWAS cohort was estimated from all post-QC SNPs in the full set of 10,000 individuals with GEMMA^30^ (Table S12).

For the YFS, we quantified the mediating effects of trait on cis-expression by separately re-estimating cis-h2g with all analyzed traits (height, BMI, TC, TG, HDL, LDL) included as fixed-effects in addition to PCs. We did not observe significant differences in any individual gene (after accounting for 3,836 genes tested) nor in the mean estimate of cis-h2g.

### Heritability estimation with summary data

As shown in ref.^54,55^, for an association study of N independent samples, the expected χ^2^ statistic is 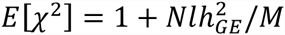, where *l* is the LD-score accounting for correlation, *M* is the number of markers, and 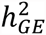 is the variance in trait explained by the imputed expression. We estimated *l* directly from the genetic values of expression to be close to independence (1.4, 1.5 for METSIM, YFS) allowing us to solve for 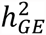 from the observed distribution of χ^2^ (or, equivalently Z^2^) statistics. We did not compute this value for the BMI GWAS because the conservative multiple GC-correction applied in that study would yield a severe downwards bias^5^.

### Evaluating prediction accuracy

Prediction accuracy was measured by five-fold cross-validation in a random sampling of 1,000 of the highly heritable genes (i.e. significant non zero cis-heritability) for each study. We evaluated three prediction schemes: i. cis-eQTL, the single most significantly associated SNP in the training set was used as the predictor; ii. the best linear predictor (BLUP)^29^, estimates the causal effect-sizes of all SNPs in the locus jointly using a single variance-component; iii. The Bayesian linear mixed model (BSLMM)^30^, which estimates the underlying effect-size distribution and then fits all SNPs in the locus jointly. For the BLUP and BSLMM, prediction was done over all post-QC SNPs using GEMMA^30^. In all instances, the R^2^ between predicted and true expression across all predicted folds was used to evaluate accuracy. On average, the cis-eQTL yielded prediction R^2^=0.08, corresponding to half of the accuracy of the best possible linear prediction (as inferred from average 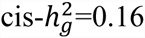 for these genes). Using all SNPs in the locus, the BLUP attained an R^2^=0.09; and the BSLMM attained an R^2^=0.10 (Methods, Fig. 2, Figure S1). The pattern was roughly the same for randomly selected genes (Figure S6). This empirical measure of accuracy deviates from theory in two ways: assuming a small number of independent SNPs at each locus, we would expect normalized BLUP accuracy 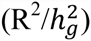 to be near 1.0 (see Equation 1 of ref. ^20^) but it is only 0.55 on average; and for a given 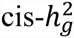, accuracy is not directly proportional to training sample size (e.g. the highest accuracy is observed in the smallest METSIM cohort, Figure S1). This suggests that data quality and population homogeneity (which differ between these cohorts) play an important role in empirical prediction accuracy.

### Imputing expression into GWAS summary statistics

Summary-based imputation was performed using the ImpG-Summary algorithm^3^ extended to train on gene expression. Let Z be a vector of standardized effect sizes (z-scores) of SNP on trait at a given cis-locus (i.e. Wald statistics 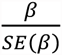). We impute the z-score of the expression and trait as a linear combination of elements of Z with weights W (these weights are precompiled from the reference panel as 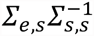 for ImpG-Summary or directly from BSLMM). *∑*_*e,s*_ is the covariance matrix between all SNPs at the locus and gene expression and *∑*_*s,s*_ is the covariance among all SNPs (i.e. linkage disequilibrium). Under null data (no association) and a multi-variate normal assumption *Z*∼ *N*(0, *∑*_*s,s*_). It follows that imputed z-score of expression and trait (*WZ*) has variance *W∑*_*s,s*_*W* ^*t*^; therefore, we use 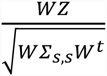 as the imputation z-score of cis-genetic effect on trait.

In practice, for each gene, all SNPs within 1Mb of the gene present in the GWAS study were selected. We computed *∑*_*s,s*_ and *∑*_*e,s*_ from the reference panel (i.e. expression and SNP data). To account for statistical noise we adjusted the diagonal of the matrix using a technique similar to ridge regression as in Pasaniuc et al^3^.

We used the YFS samples that were assayed for SNPs, phenotype, and expression to assess the consistency of individual-level and summary-based TWAS. We first computed GWAS association statistics between phenotype (height) and SNP and used them in conjunction with the expression data to impute summary-based TWAS statistics. The TWAS statistics were compared to those from the simple regression of (height ∼ expression) in the YFS data. We observed a correlation of 0.415 (Supplementary Figure S13), consistent with an average cis-h2g of 0.17 (≈0.415^2) observed for these genes. When restricting to a regression of (height ∼ cis component of expression) we observed a correlation of 0.998 to the summary based TWAS, demonstrating the equivalence of the two approaches when using in-sample LD.

For imputation using an external cis-eQTL study, *∑*_*e,s*_estimated from the available cis-eQTL association statistics instead of directly in the training data. For the MuTHER SNPs, this was estimated by computing the corresponding cis-eQTL T-statistic; solving for R^2^=T^2^/(T^2^+N-2) where N was the study size; and converting back to signed *r* using the effect-size direction. The LD matrix was estimated from the METSIM as before. In the case where the LD matrix matches that of the eQTL study, this approach is mathematically identical to training on individual-level data. Otherwise, differences in LD will introduce noise which is expected to be unbiased assuming no relationship between these differences and eQTL effect-size. These out-of-sample expression effect-sizes from the MuTHER study allowed us to evaluate the impact of the LD-reference panel size on accuracy. We compared predictors trained using an LD-reference panel from the ∼600 METSIM expression samples to those trained on the 6,000 unrelated METSIM GWAS individuals and found highly significant consistency (*ρ*=0.97; Figure S7) with slight Z-score inflation in the smaller panel.

### Power analysis of summary-based method

Simulations to evaluate the summary-based method were performed in 6,000 unrelated METSIM GWAS individuals. 100 genes and the SNPs in the surrounding 1MB were randomly selected for testing. For each gene, normally distributed gene expression was simulated as *E* = **X***β*+ ε; where **X** is a matrix of the desired number of causal genotypes, sampled randomly from the locus; *β* is a vector of normally distributed effect-sizes for each causal variant; and *ϵ* is a vector of normally distributed noise to achieve a cis-h2g of 0.17 (corresponding to the mean observed in our significant gene sets). 1,000 individuals with SNPs and simulated expression were then withheld for training the predictors. For the remaining 5,000 individuals, normally distributed noise was applied to the expression to generate a heritable phenotype where expression explained 0.10/180 or 0.20/180 of the phenotypic variance (the former corresponding to the average effect-sizes for associated genes observed in a large GWAS of height^56^ and the latter to high-effect loci). Association between SNP and phenotype was estimated in the 5,000 individuals (standard Z-score), and the phenotype generation repeated with different environmental noise (up to 60 times) to generate results from multiple GWAS sub-studies. Association statistics from each run were then meta-analyzed to reach precision corresponding to a larger GWAS of desired size (up to 300,000).

The above procedure is equivalent to assuming that LD and MAF do not change across sub-studies, and we believe these assumptions are reasonable for large studies of a homogenous population where individual-level data is not used. To verify this assumption, we re-ran the main set of simulations using the HAPGEN2 algorithm^57^ to simulate new genotypes for each sub-study from a phased reference panel of the 5,000 held-out samples. This method models population demography and uses a haplotype-copying model to generate new individuals based on a phased reference panel. Under this complex model, none of the results were substantially different from the previous simulations (Supplementary Figure S12), and so we used the phenotype regeneration procedure due to its computational efficiency.

Detecting a locus was defined as follows. The single most significant trait associated SNP was taken as the GWAS association, considered detected if GWAS significance was <5×10^−8^. The single most significant eQTL in the training set was taken as the eQTL-guided association (eGWAS), and considered detected if GWAS significance was <0.05/15,000. The TWAS association was measured by training the imputation algorithm on the 1,000 held-out samples with expression and imputing into the GWAS summary statistics, and considered detected if significance was <0.05/15,000. The entire procedure was repeated 500 times (5 per gene) and power was estimated by counting the fraction of instances where each method detected the locus. As in the cross-validation analysis, training on the genetic component of expression instead of the overall expression consistently increased TWAS power by ∼10% (Figure S8). Two null expression models were tested by generating gene expression for the 1,000 held-out samples that was standard normal as well as heritable expression (cis-h2g=0.17) with GWAS Z-scores drawn from the standard normal (Table S2).

Lastly, we evaluated the confounded model where expression and trait had the same causal variants but independent effect-sizes (Figure 2G). The case of a single causal variant with independent effects is statistically indistinguishable from a true causal model. Consider the following generative model for expression (*E*) where *x* is the causal variant, *β* is the effect, and *ϵ* is the scaled environmental effect:

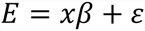

for two possible models for phenotype (Y):

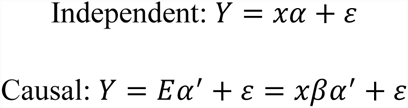

the models will be identical if *α* = *βα’* and therefore cannot be distinguished by a statistical test of *α’* being non-zero without direct mediation analysis. Similar intuition applies when multiple causal variants or tags are present, where power is expected to decrease with the *r*^2^ between the true and observed effect^58^. Consistent with theory, our simulations show that the two models are equally likely to be detected by all methods (Figure S9, SEEE). In the case of multiple causal variants, the detection rate of the independent scenario is much lower and roughly equal for TWAS or eGWAS (Figure S9).

### Power comparison to COLOC

We used a similar simulation framework as above to compare TWAS to the recently proposed COLOC method. COLOC uses summary data from eQTL and GWAS studies and a Bayesian framework to identify the subset of GWAS signals that co-localize with eQTLs. Because COLOC relies on priors of association to produce posterior probabilities of co-localization, we sought to identify a significance threshold that would make a fair comparison to the TWAS p-value-based threshold. Specifically, we ran both methods on a realistic null expression simulation (with the generative model described previously): the expression was sampled from a null standard normal for 1,000 individuals and eQTLs computed; the trait associations were derived from a simulated 300,000 GWAS with a single typed causal variant that explained 0.001 variance of the trait (high effect). We believe this scenario is both realistic and consistent with the GWAS assumptions of COLOC. We then empirically identified the statistical threshold for COLOC and TWAS that would yield a 5% false discovery rate: co-localization statistic PPA > 0.17 for COLOC, and P<0.05 for TWAS. We note that this empirical COLOC threshold is much less stringent than PPA>0.8 used in the COLOC paper (PPA>0.8 would yield lower power for COLOC in our simulations). These thresholds were subsequently used as cutoffs to evaluate the power to detect an expression-trait association in simulations with a true effect (Supplementary Figure S11, S14). The reported power is for a single locus and we did not attempt to quantify genome/transcriptome-wide significance.

### Individual-level analysis of METSIM GWAS

We imputed the significantly heritable genes into the METSIM GWAS cohort of 5,500 unrelated individuals with individual-level genotypes (and unmeasured expression). We then tested the imputed expression for obesity-related traits: body mass index (BMI); triglycerides (TG); waist-hip-ratio (WHR); and fasting insulin levels (INS). Overall, the evaluated traits exhibited high phenotypic and genetic correlation as well as highly significant genome-wide 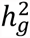 ranging from 23-36% (Table S12) consistent with common variants having a major contribution to disease risk^7^. Association was assessed using standard regression as well as a mixed-model that accounted for relatedness and phenotypic correlation^30^ with similar results. The effective number of tests for each trait was estimated by permuting the phenotypes 10,000 times and, for each permutation, re-running the association analysis on all predicted genes. For each trait P_*perm*_, the P-value in the lowest 0.05 of the distribution, was computed and the effective number of tests was 0.05/P_*perm*_, reported in Table S13. All phenotypes were shuffled together, so any phenotypic correlation was preserved. The effective number of tests corresponded to 88-95% of the total number of genes, indicating a small amount of statistical redundancy.

After accounting for multiple testing in each trait, six loci were significant (Table S14); five of which were confirmed by genome-wide significant SNP associations in this cohort or in larger studies. The best cis-eQTL in each locus was less significantly associated than the imputed expression in 5/6 loci, further underscoring the increased power of the TWAS approach. The TWAS identified one novel gene that had not been previously observed in this or published GWAS: ENO3 associated with TG and fasting insulin (INS). We investigated this gene further in the METSIM samples with both phenotypes and expression, and found a nominal association between the genetic value of expression and INS at P=0.02, explaining 1.8% of trait variance (with phenotypes which had not been used to identify the initial association). This association was primarily driven by the top eQTL (rs9914087, P=8×10^−09^ for eQTL, P=0.03 for association to INS). Though validation in a larger cohort is needed, this initial result supports a link between ENO3 expression and fasting insulin in this population.

To evaluate the TWAS approach, we computed phenotype association statistics for the 5,500 unrelated individuals and re-ran the analysis using only these summary statistics and the same expression reference panels. The resulting TWAS associations were nearly identical to the direct TWAS associations across the four traits (Pearson *ρ*=0.96). Reassuringly, the TWAS was generally more conservative than the direct estimates (Figure S5).

### Refining trait-associated genes at known loci

We first sought to validate the performance of TWAS in identifying trait-effecting genes at previously implicated loci. We focused on GWAS data from a recent large study of height^6^ that identified 697 genome-wide significant variants in 423 loci, and conducted the summary-based TWAS over all genes in these loci using YFS as training data. If the TWAS was identifying trait-effecting genes, we would expect the expression of these genes to be associated with height in an independent sample. Because the YFS individuals had been phenotyped for height and not tested in the GWAS, we could evaluate this directly. At each locus, we considered three strategies for selecting a single causal gene: 1) the gene nearest to the most significantly associated SNP; 2) the gene for which the index SNP is the strongest eQTL in the training data; 3) the most significant TWAS gene. For each strategy, we then constructed a risk-score using the genetic value of expression for the selected gene weighted by the corresponding TWAS Z-score (see Methods). The R^2^ between the risk score and the height phenotype was 0.038 (nearest); 0.031 (eQTL); and 0.054 (TWAS); with TWAS significantly higher than the others (Table S3). Further restricting to the 263 loci where the TWAS gene was not the nearest, the R^2^ was 0.011 (nearest) and 0.032 (TWAS); with only the latter significant in a joint model (P=7×10^−3^). We separately used GCTA to estimate the heritability of height explained by all of the genes selected by each algorithm by constructing a GRM from the selected genes. In contrast to the risk score, this does not assume pre-defined weights on each genes but allows them to be fit by the REML model. Results were comparable, with only the TWAS-selected genes explaining significantly non-zero heritability (Table S3).

### Validation analysis in lipid GWAS data

We evaluated the performance of TWAS by identifying significantly associated genes in the 2010 lipid study that did not overlap a genome-wide significant SNP, and looking for newly genome-wide significant SNPs in the expanded 2013 study. The P-value for the number of genes with increased significance and genome-wide significance in the 2013 study was computed by a hypergeometric test, with background probabilities estimated from the set of significantly heritable genes. Of the genes not overlapping a significant locus in the 2010 study, 70% had a more significant SNP in the 2013 study and 3.5% overlapped a genome-wide significant SNP (P<5×10^−08^).

### Meta-analysis of imputed expression from multiple tissues

We proposed a novel omnibus test for significant association across predictions from all three cohorts. Because the imputation is made into the same GWAS cohort, correlation between predictors must be accounted for. For each gene *i*, we estimated a correlation matrix **C**_*i*_ by predicting from the three tissues into the ∼5,500 unrelated METSIM GWAS individuals (though any large panel from the study population could be used). This correlation includes both the genetic correlation of expression as well as any correlated error in the predictors, thus capturing all redundancy. On average, a correlation of 0.01, 0.01, and 0.43 was observed between YFS:METSIM, NTR:METSIM, and YFS:NTR, highlighting the same tissue of origin the last pair. We then used the three-entry vector of TWAS predictions, Z_*i*_, to compute the statistic 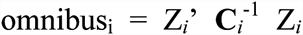 which is approximately χ^2^ (3-dof) distributed and provides an omnibus test for effect in any tissue while accounting for correlation^59,60^. Though the correlation observed in our data was almost entirely driven by the YFS:NTR blood datasets, we expect this to be an especially useful strategy for future studies with many correlated tissues. An alternative approach would be to perform traditional meta-analysis across the three cohorts and then predict the TWAS effect. However, this would lose power when true eQTL effect-sizes (or LD) differ across the cohorts, which we have empirically observed to be the case looking at predictor correlations above. The proposed omnibus test aggregates different effects across the studies, at the cost of additional degrees of freedom.

### Gene permutation test

We conducted a permutation test to quantify the TWAS contribution conditional on the observed trait effects. For each gene, the expression labels were randomly shuffled and the summary-based TWAS analysis trained on the resulting expression to compute a permuted expression-trait Z-score. This procedure was repeated 1,000 times for every locus to establish a null expression distribution, and a p-value for the real expression Z-score was computed by Z-test against this null. Because the GWAS statistics were unchanged, this procedure computes a distribution for (trait ∼ SNP + GE) where GE is null but the true SNP effect is retained, testing whether the GE contribution is individually significant beyond the contribution of the SNP. For example, consider a locus with a single highly significant GWAS association *Z*_*i*_, and expression weights trained in N individuals: the resultant TWAS weights would be normally distributed with variance=1/N, and the TWAS Z-score would be normally distributed with 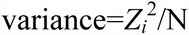, which may yield inflated estimates when 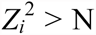 or when both distributions have non-zero means. The permutation evaluates this null while accounting for the true correlation between markers in the locus. Note that failing the permutation test only indicates lack of power to show that the expression significantly refines the direct SNP-trait signal.

We again sought to validate this test by focusing on known loci in the height GWAS and the YFS cohort with height measured independently (see above). Using YFS expression as training, we used the TWAS approach in the independent height GWAS data to identify 181 significant genes that overlapped previously known height loci, of which 33 passed the permutation test. These 33 genes had evidence of a significant contribution to trait beyond the SNP-trait effects at the locus, indicative of allelic heterogeneity at these loci. As before, we constructed a risk score using the genetic value of these 33 genes weighted by their Z-score in the TWAS; and a standard genetic risk score^61^ using the best GWAS SNP in each locus. The two scores were evaluated for association to the true height phenotype in the YFS, yielding an R^2^ of 0.008 (best SNP) and 0.016 (TWAS gene) respectively (Table S3). In a joint regression with both scores, only the TWAS score was significant (P=2×10^−3^). This confirms that TWAS predictions which remain significant after permutation are more strongly associated with phenotype than the single best SNP at the locus.

### Relationship to genetic covariance

Our tests relate to previously defined estimators of genetic correlation and covariance between traits. We consider two definitions of genetic covariance at a locus: 1) the covariance between the genetic component of expression and the genetic component of trait; 2) the covariance between the causal effect sizes for expression and the causal effect-sizes for trait. Under assumptions of independent effect-sizes, these definitions yield asymptotically identical quantities^62^. Assuming a substantially large training set where the genetic component of expression can be perfectly predicted, the direct TWAS tests for a significant association between the genetic component of expression and the trait; equivalent to testing definition #1 for a polygenic trait. Likewise, the TWAS tests for a significant sum of products of the causal expression effect sizes and the causal trait effect sizes; equivalent to definition #2 up to a scaling factor. The TWAS approach therefore fits naturally with the broader study of genetic of multiple phenotypes.

## Acknowledgments

We thank the individuals who participated in the study. We also acknowledge Liming Yang for helpful discussions that have improved the quality of this manuscript. We would also like to thank Karen Mohlke, Michael Boehnke and Francis Collins for help with the METSIM data. This work was funded in part by NIH grants F32 GM106584 (S.G.), R1 GM053725 (B.P.), R01 GM105857 (A.L.P., S.G., G.B.), HL-28481 (P.P., A.J.L., M.C.), HL-095056 (P.P., B.P.), and by the NIH training grant in Genomic Analysis and Interpretation T32 HG002536 (A.K.)

---

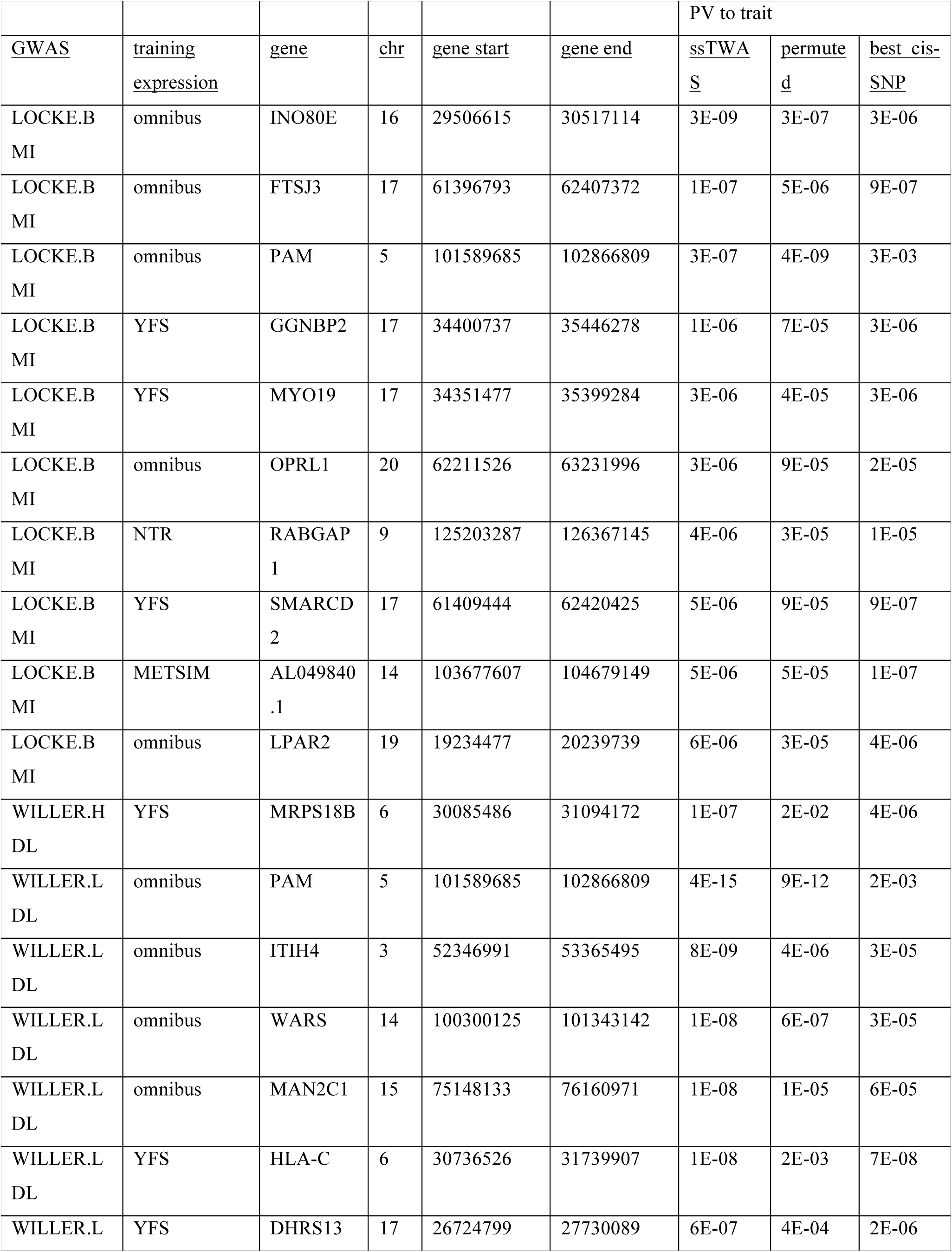
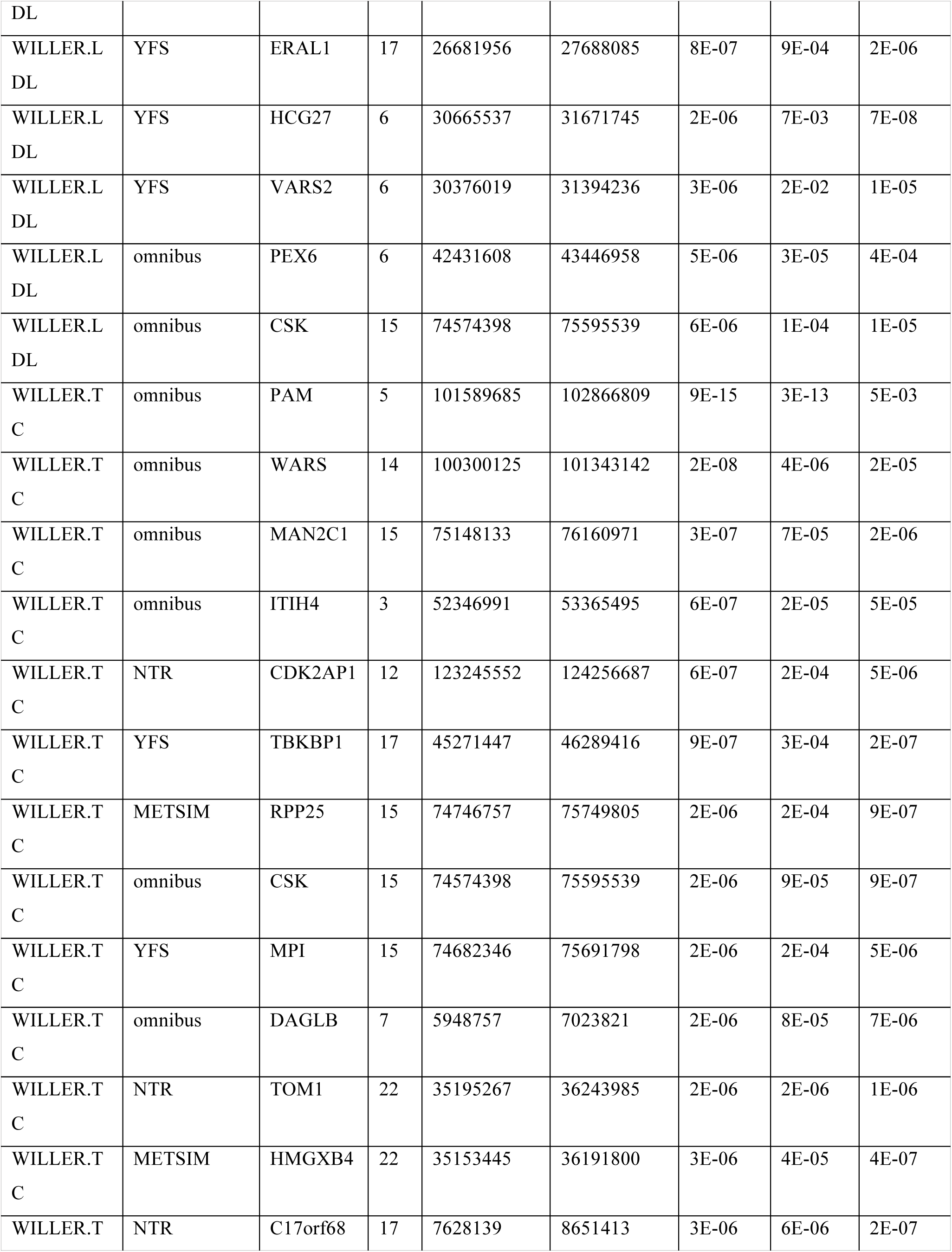
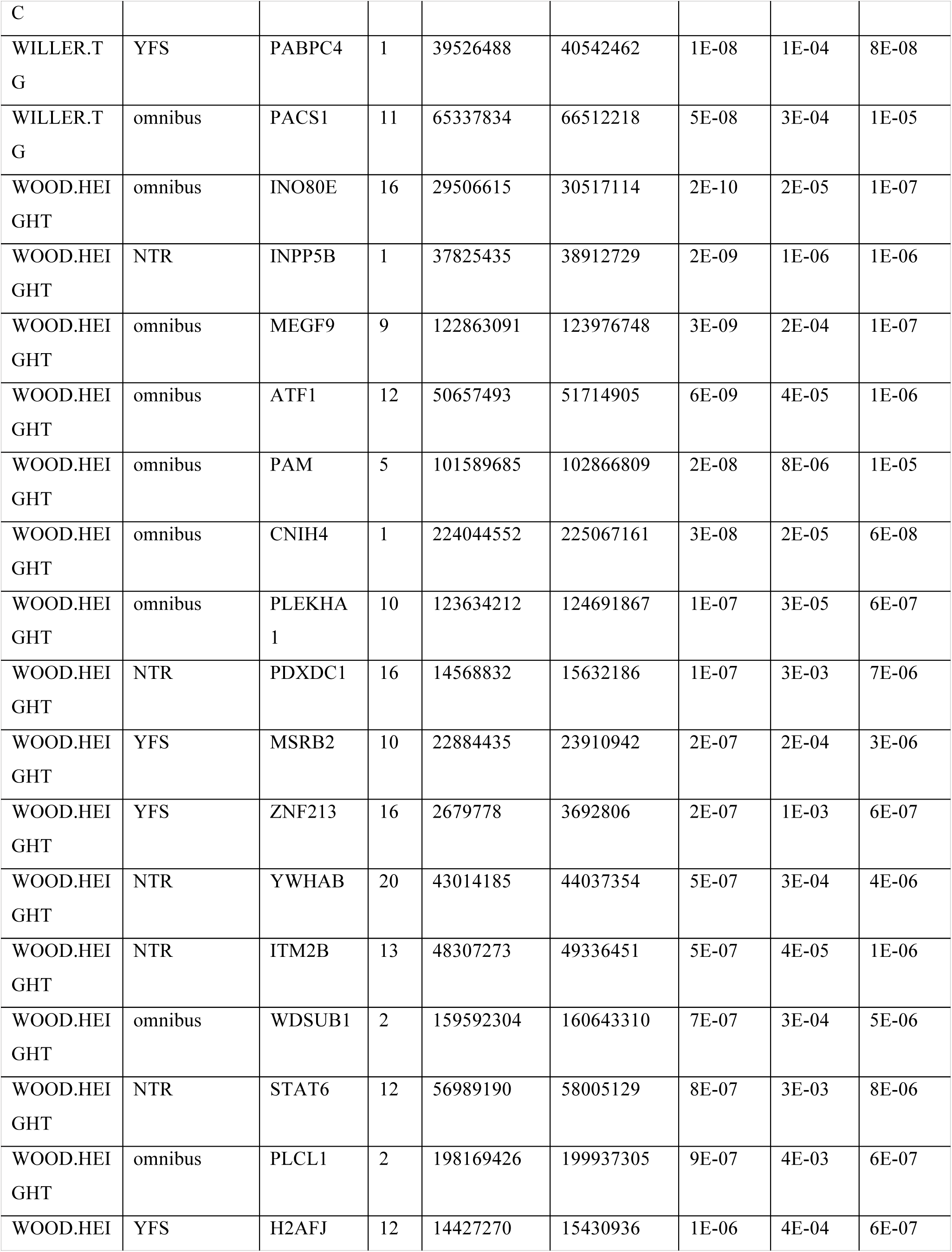
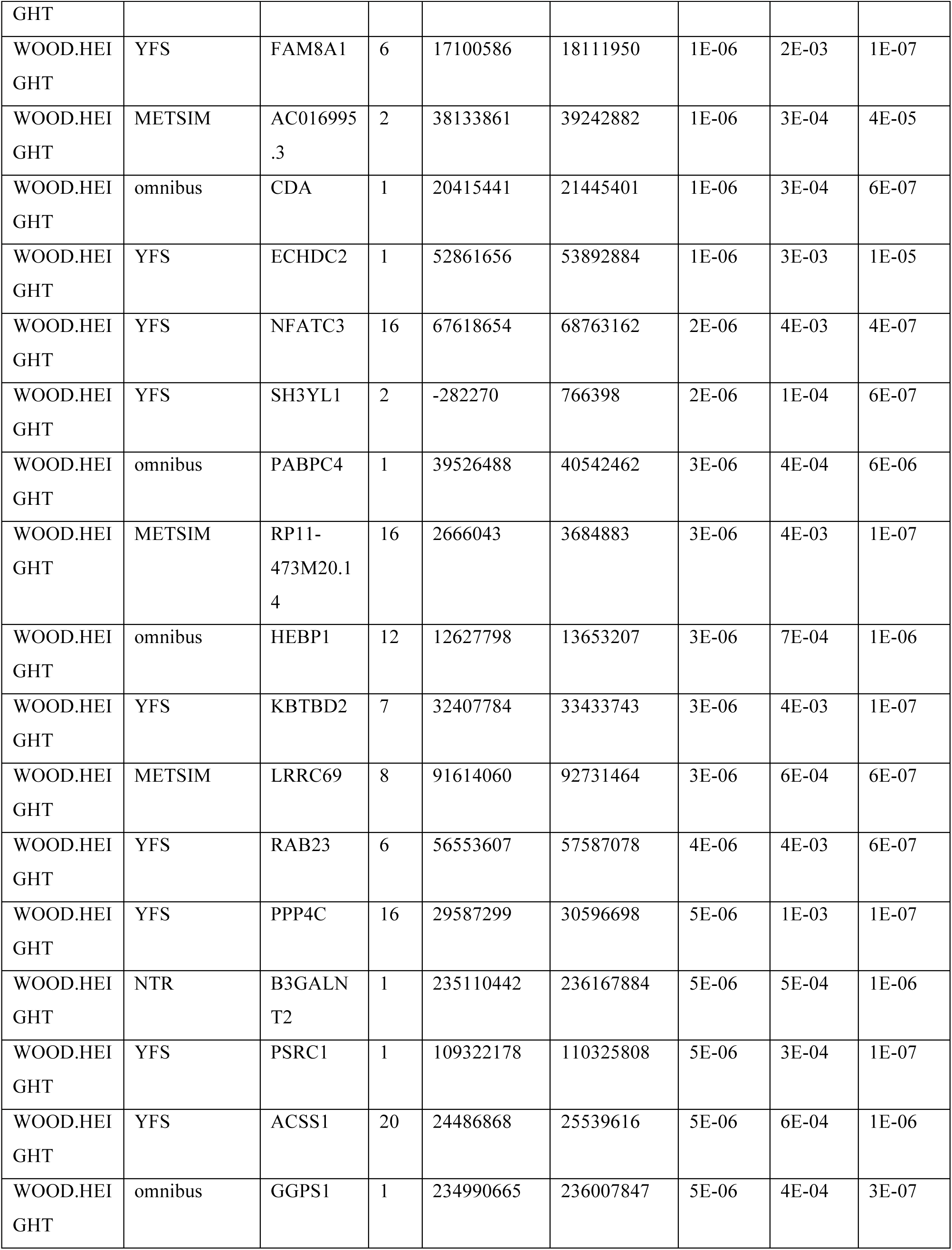

## References

1. Yang, J. et al. Conditional and joint multiple-SNP analysis of GWAS summary statistics identifies additional variants influencing complex traits. Nature genetics 44, 369–375, S361-363, doi:10.1038/ng.2213 (2012).

2. Lee, D., Bigdeli, T. B., Riley, B. P., Fanous, A. H. & Bacanu, S. A. DIST: direct imputation of summary statistics for unmeasured SNPs. Bioinformatics 29, 2925–2927, doi:10.1093/bioinformatics/btt500 (2013).

3. Pasaniuc, B. et al. Fast and accurate imputation of summary statistics enhances evidence of functional enrichment. Bioinformatics 30, 2906–2914, doi:10.1093/bioinformatics/btu416 (2014).

4. Global Lipids Genetics, C. et al. Discovery and refinement of loci associated with lipid levels. Nature genetics 45, 1274–1283, doi:10.1038/ng.2797 (2013).

5. Locke, A. E. et al. Genetic studies of body mass index yield new insights for obesity biology. Nature 518, 197–206, doi:10.1038/nature14177 (2015).

6. Wood, A. R. et al. Defining the role of common variation in the genomic and biological architecture of adult human height. Nature genetics 46, 1173–1186, doi:10.1038/ng.3097 (2014).

7. Visscher, P. M., Brown, M. A., McCarthy, M. I. & Yang, J. Five years of GWAS discovery. American journal of human genetics 90, 7–24, doi:10.1016/j.ajhg.2011.11.029 (2012).

8. Musunuru, K. et al. From noncoding variant to phenotype via SORT1 at the 1p13 cholesterol locus. Nature 466, 714–719, doi:10.1038/nature09266 (2010).

9. Lappalainen, T. et al. Transcriptome and genome sequencing uncovers functional variation in humans. Nature 501, 506–511, doi:10.1038/nature12531 (2013).

10. Zhang, X. et al. Identification of common genetic variants controlling transcript isoform variation in human whole blood. Nature genetics, doi:10.1038/ng.3220 (2015).

11. Westra, H. J. et al. Systematic identification of trans eQTLs as putative drivers of known disease associations. Nature genetics 45, 1238–1243, doi:10.1038/ng.2756 (2013).

12. Albert, F. W. & Kruglyak, L. The role of regulatory variation in complex traits and disease. Nature reviews. Genetics, doi:10.1038/nrg3891 (2015).

13. Raj, T. et al. Polarization of the effects of autoimmune and neurodegenerative risk alleles in leukocytes. Science 344, 519–523, doi:10.1126/science.1249547 (2014).

14. Letourneau, A. et al. Domains of genome-wide gene expression dysregulation in Down’s syndrome. Nature 508, 345–350, doi:10.1038/nature13200 (2014).

15. Davis, L. K. et al. Partitioning the heritability of Tourette syndrome and obsessive compulsive disorder reveals differences in genetic architecture. PLoS genetics 9, e1003864, doi:10.1371/journal.pgen.1003864 (2013).

16. Teslovich, T. M. et al. Biological, clinical and population relevance of 95 loci for blood lipids. Nature 466, 707–713, doi:10.1038/nature09270 (2010).

17. Yang, J. et al. Common SNPs explain a large proportion of the heritability for human height. Nature genetics 42, 565–569, doi:10.1038/ng.608 (2010).

18. Yang, J., Lee, S. H., Goddard, M. E. & Visscher, P. M. GCTA: a tool for genome-wide complex trait analysis. American journal of human genetics 88, 76–82, doi:10.1016/j.ajhg.2010.11.011 (2011).

19. Gusev, A. et al. Partitioning heritability of regulatory and cell-type-specific variants across 11 common diseases. American journal of human genetics 95, 535–552, doi:10.1016/j.ajhg.2014.10.004 (2014).

20. Wray, N. R. et al. Pitfalls of predicting complex traits from SNPs. Nature reviews. Genetics 14, 507–515, doi:10.1038/nrg3457 (2013).

21. Nuotio, J. et al. Cardiovascular risk factors in 2011 and secular trends since 2007: the Cardiovascular Risk in Young Finns Study. Scandinavian journal of public health 42, 563–571, doi:10.1177/1403494814541597 (2014).

22. Raitakari, O. T. et al. Cohort profile: the cardiovascular risk in Young Finns Study. International journal of epidemiology 37, 1220–1226, doi:10.1093/ije/dym225 (2008).

23. Wright, F. A. et al. Heritability and genomics of gene expression in peripheral blood. Nature genetics 46, 430–437, doi:10.1038/ng.2951 (2014).

24. Grundberg, E. et al. Mapping cis- and trans-regulatory effects across multiple tissues in twins. Nature genetics 44, 1084–1089, doi:10.1038/ng.2394 (2012).

25. Nicolae, D. L. et al. Trait-associated SNPs are more likely to be eQTLs: annotation to enhance discovery from GWAS. PLoS genetics 6, e1000888, doi:10.1371/journal.pgen.1000888 (2010).

26. Torres, J. M. et al. Cross-tissue and tissue-specific eQTLs: partitioning the heritability of a complex trait. American journal of human genetics 95, 521–534, doi:10.1016/j.ajhg.2014.10.001 (2014).

27. Buil, A. et al. Gene-gene and gene-environment interactions detected by transcriptome sequence analysis in twins. Nature genetics 47, 88–91, doi:10.1038/ng.3162 (2015).

28. Nica, A. C. et al. Candidate causal regulatory effects by integration of expression QTLs with complex trait genetic associations. PLoS genetics 6, e1000895, doi:10.1371/journal.pgen.1000895 (2010).

29. Robinson, G. K. That BLUP is a good thing: the estimation of random effects. Statistical science, 15–32 (1991).

30. Zhou, X., Carbonetto, P. & Stephens, M. Polygenic modeling with bayesian sparse linear mixed models. PLoS genetics 9, e1003264, doi:10.1371/journal.pgen.1003264 (2013).

31. Dudbridge, F. Power and predictive accuracy of polygenic risk scores. PLoS genetics 9, e1003348, doi:10.1371/journal.pgen.1003348 (2013).

32. Chatterjee, N. et al. Projecting the performance of risk prediction based on polygenic analyses of genome-wide association studies. Nature genetics 45, 400–405, 405e401-403, doi:10.1038/ng.2579 (2013).

33. Brown, C. D., Mangravite, L. M. & Engelhardt, B. E. Integrative modeling of eQTLs and cis-regulatory elements suggests mechanisms underlying cell type specificity of eQTLs. PLoS genetics 9, e1003649, doi:10.1371/journal.pgen.1003649 (2013).

34. Wen, X., Luca, F. & Pique-Regi, R. Cross-population joint analysis of eQTLs: fine mapping and functional annotation. PLoS genetics 11, e1005176, doi:10.1371/journal.pgen.1005176 (2015).

35. Wood, A. R. et al. Another explanation for apparent epistasis. Nature 514, E3–5, doi:10.1038/nature13691 (2014).

36. Giambartolomei, C. et al. Bayesian test for colocalisation between pairs of genetic association studies using summary statistics. PLoS genetics 10, e1004383, doi:10.1371/journal.pgen.1004383 (2014).

37. Pritchard, J. K. & Cox, N. J. The allelic architecture of human disease genes: common disease-common variant…or not? Human molecular genetics 11, 2417–2423 (2002).

38. Wen, W. et al. Meta-analysis of genome-wide association studies in East Asian-ancestry populations identifies four new loci for body mass index. Human molecular genetics 23, 5492–5504, doi:10.1093/hmg/ddu248 (2014).

39. Pers, T. H. et al. Biological interpretation of genome-wide association studies using predicted gene functions. Nature communications 6, 5890, doi:10.1038/ncomms6890 (2015).

40. Smith, G. D. & Ebrahim, S. ’Mendelian randomization’: can genetic epidemiology contribute to understanding environmental determinants of disease? International journal of epidemiology 32, 1–22 (2003).

41. Pickrell, J. Fulfilling the promise of Mendelian randomization. (2015).

42. Pierce, B. L. & Burgess, S. Efficient design for Mendelian randomization studies: subsample and 2-sample instrumental variable estimators. American journal of epidemiology 178, 1177–1184, doi:10.1093/aje/kwt084 (2013).

43. Gusev, A. et al. Quantifying missing heritability at known GWAS loci. PLoS genetics 9, e1003993, doi:10.1371/journal.pgen.1003993 (2013).

44. Dimas, A. S. et al. Common regulatory variation impacts gene expression in a cell type-dependent manner. Science 325, 1246–1250, doi:10.1126/science.1174148 (2009).

45. Montgomery, S. B. et al. Transcriptome genetics using second generation sequencing in a Caucasian population. Nature 464, 773–777, doi:10.1038/nature08903 (2010).

46. Pickrell, J. K. Joint analysis of functional genomic data and genome-wide association studies of 18 human traits. American journal of human genetics 94, 559–573, doi:10.1016/j.ajhg.2014.03.004 (2014).

47. Kichaev, G. et al. Integrating functional data to prioritize causal variants in statistical fine-mapping studies. PLoS genetics 10, e1004722, doi:10.1371/journal.pgen.1004722 (2014).

48. Stancakova, A. et al. Hyperglycemia and a common variant of GCKR are associated with the levels of eight amino acids in 9,369 Finnish men. Diabetes 61, 1895–1902, doi:10.2337/db11-1378 (2012).

49. Stancakova, A. et al. Changes in insulin sensitivity and insulin release in relation to glycemia and glucose tolerance in 6,414 Finnish men. Diabetes 58, 1212–1221, doi:10.2337/db08-1607 (2009).

50. Turchin, M. C. et al. Evidence of widespread selection on standing variation in Europe at height-associated SNPs. Nature genetics 44, 1015–1019, doi:10.1038/ng.2368 (2012).

51. Dobin, A. et al. STAR: ultrafast universal RNA-seq aligner. Bioinformatics 29, 15–21, doi:10.1093/bioinformatics/bts635 (2013).

52. Stegle, O., Parts, L., Piipari, M., Winn, J. & Durbin, R. Using probabilistic estimation of expression residuals (PEER) to obtain increased power and interpretability of gene expression analyses. Nature protocols 7, 500–507, doi:10.1038/nprot.2011.457 (2012).

53. Shi, W., Oshlack, A. & Smyth, G. K. Optimizing the noise versus bias trade-off for Illumina whole genome expression BeadChips. Nucleic acids research 38, e204, doi:10.1093/nar/gkq871 (2010).

54. Finucane, H. K. et al. Partitioning heritability by functional category using GWAS summary statistics. (2015).

55. B. K. et al. LD Score regression distinguishes confounding from polygenicity in genome-wide association studies. Nature genetics 47, 291–295, doi:10.1038/ng.3211 (2015).

56. Lango Allen, H. et al. Hundreds of variants clustered in genomic loci and biological pathways affect human height. Nature 467, 832–838, doi:10.1038/nature09410 (2010).

57. Su, Z., Marchini, J. & Donnelly, P. HAPGEN2: simulation of multiple disease SNPs. Bioinformatics 27, 2304–2305, doi:10.1093/bioinformatics/btr341 (2011).

58. Pritchard, J. K. & Przeworski, M. Linkage disequilibrium in humans: models and data. American journal of human genetics 69, 1–14, doi:10.1086/321275 (2001).

59. Bolormaa, S. et al. A multi-trait, meta-analysis for detecting pleiotropic polymorphisms for stature, fatness and reproduction in beef cattle. PLoS genetics 10, e1004198, doi:10.1371/journal.pgen.1004198 (2014).

60. Zhu, X. et al. Meta-analysis of correlated traits via summary statistics from GWASs with an application in hypertension. American journal of human genetics 96, 21–36, doi:10.1016/j.ajhg.2014.11.011 (2015).

61. International Schizophrenia, C. et al. Common polygenic variation contributes to risk of schizophrenia and bipolar disorder. Nature 460, 748–752, doi:10.1038/nature08185 (2009).

62. Bulik-Sullivan, B. et al. An Atlas of Genetic Correlations across Human Diseases and Traits. (2015).

